# Self-pruning in tree crowns is influenced by functional strategies and neighborhood interactions

**DOI:** 10.1101/2024.10.17.618957

**Authors:** Shan Kothari, Jon Urgoiti, Christian Messier, William S. Keeton, Alain Paquette

## Abstract

As canopy closure causes forest stands to face increasing light limitation, trees’ lower branches begin to die back. This process, called self-pruning, defines a crown’s base and depth and shapes the structure of entire stands. Self-pruning is often thought to occur after shading causes individual branches to transition from net carbon sources to sinks. Under this explanation, we would expect resource-conservative and shade-tolerant species to initiate self-pruning under deeper shade because their branches need less light to maintain a positive carbon balance. However, the notion that branches are fully autonomous may be complicated by ‘correlative inhibition,’ in which plants preferentially allocate resources towards sunlit branches. Consistent with this idea, we predicted that within species, trees with sunlit tops would initiate self-pruning at a higher light threshold. Lastly, we predicted that plot-level diversity in self-pruning strategies would correlate with productivity and total crown volume. We tested these predictions in an experiment where 12 temperate tree species were planted in plots of varying diversity and composition. We measured characteristics of crown size and position as well as the light level at the crown base (denoted *L_base_*), which we took as an estimate of the light threshold of self-pruning. As we predicted, more shade-tolerant and resource-conservative species self-pruned at a deeper level of shade (lower *L_base_*). In addition, most species had higher *L_base_* when they had more light at the crown top, suggestive of correlative inhibition. With respect to their neighbors’ traits, though, conservative and acquisitive species showed contrary patterns of plasticity: conservative species had lower *L_base_* around conservative neighbors, while acquisitive species had lower *L_base_* around acquisitive neighbors. However, all species declined in crown depth when they grew alongside larger, more acquisitive neighbors. As predicted, plots with a greater interspecific diversity of *L_base_* had greater basal area and crown volume. Using simulations, we showed that plasticity in crown depth between monocultures and mixtures strengthened the relationship with crown volume, primarily due to competitive release experienced by acquisitive species. By placing shade-induced self-pruning in a comparative context, we clarify how forest function emerges from competition for light between individual trees.

## Introduction

The arrangement of tree crowns in three-dimensional space determines how much light they can capture, thereby exerting a strong influence on forest productivity and development. The shape and size of these crowns is influenced by the process of self-pruning—the regulated loss of branches. Self-pruning in the lower parts of a tree crown defines the live crown base, and thus (along with tree height) determines crown depth (Mäkelä 1997). As a result, self-pruning may play an important part in defining the light acquisition strategies of species and creating the complex, multi-layered vertical structure of light-limited forests. Self-pruning also matters from a silvicultural perspective: since knots left by branches’ attachments to the trunk are a main cause of reductions in lumber grade, the pruning and natural occlusion of knots over time often increase the value of wood (Trincado & Burkhart 2009). As a result, foresters often aim to choose species, environments, or planting conditions that reduce branching and increase self-pruning rates without compromising stand growth (Mäkinen 2002; Garber et al. 2008; Kint et al. 2010).

However, there is little comparative research on self-pruning and its role in trees’ strategies under competition for light. Self-pruning tends to occur in older branches of the lower crown as they become more and more shaded during stand development. Researchers have often conjectured or assumed that individual branches are pruned as diminishing access to light causes them to transition from net carbon sources to sinks (Prentice & Leemans 1990; Sorrensen-Cothern et al. 1993)—an idea that has received some empirical support (Witowski 1997). For simplicity, branches are often taken to be autonomous—they do not rely on carbon from other branches, and they live or die regardless of the carbon status of the whole plant (Sprugel et al. 1991). However, some studies have challenged the idea of branch autonomy (Henriksson 2001; Sprugel 2002; Schoonmaker et al. 2014). For example, trees whose upper crowns are well-lit sometimes prune their lower branches at a higher level of light than trees that are fully shaded, an instance of ‘correlative inhibition’ (Novoplansky et al. 1989; Stoll & Schmid 1998; Takenaka 2000; Sprugel 2002). This phenomenon may result from preferential allocation of water and nutrients to branches that assimilate the most carbon, which may cause photosynthetic decline or hydraulic failure in shaded lower branches (Protz et al. 2000). These findings indicate that self-pruning is more complex than complete branch autonomy would imply, but remain consistent with the perspective that it results from declines in branch carbon balance through shading.

Under a carbon balance-based explanation, shade-tolerant species may be expected to self-prune at a lower level of light. This hypothesis is based on the assumption that shade-tolerant species have a lower light compensation point (LCP) than shade-intolerant species, meaning that the amount of light required for their carbon balance to break even is lower (Givnish 1988; Craine & Reich 2005; Lusk & Jorgensen 2013). (In this case, the relevant definition of the LCP may include respiration not just by the branch and its leaves, but also by any other living tissues that would no longer be needed if the branch were shed—an idea closely related to the ‘effective light compensation point’ of Givnish et al. [1988].) The idea that late-successional, shade-tolerant species require deeper shade to initiate self-pruning may help explain why they have longer crowns with lower bases (Poorter et al. 2006) and themselves cast a deeper shade (Zeide 1985; Canham et al. 1994). Like shade tolerance, conservative leaf-economic traits (*sensu* Wright et al. 2004) may be expected to reduce the LCP (Baltzer & Thomas 2007a, b; Baltzer & Thomas 2010; Falster et al. 2018), causing self-pruning to be initiated at a lower light level.

Variation in self-pruning strategies could contribute to the emergent canopy structure of diverse forests. More generally, variation in crown architecture and position may allow forests to achieve greater: (1) crown complementarity, such that crowns overlap less in space (Williams et al. 2017); and (2) canopy packing, such that there is more total crown volume per unit ground area (Pretzsch 2014; Jucker et al. 2015). These phenomena may allow the community to capture the available light more completely and use it more efficiently (Sapijanskas et al. 2014; Pretzsch 2014; Williams et al. 2017; Rissanen et al. 2019; Duarte et al. 2021). Once the canopy closes and light becomes a limiting resource, canopy packing and complementarity are likely to help explain the commonly observed pattern that diversity has a positive influence on productivity across a variety of forested ecosystems (Paquette & Messier 2011; Huang et al. 2018; Urgoiti et al. 2022; Feng et al. 2022).

Past studies have noted that crown complementarity (Williams et al. 2017) and canopy packing (Jucker et al. 2015) are greater in mixtures that vary in shade tolerance. These effects arise from a combination of vertical stratification and differences in crown shape, due to both interspecific variation and plasticity. If commonly measured functional traits and shade tolerance are related to crown architecture—via their influence on self-pruning or otherwise—it may help explain why both functional diversity (Bongers et al. 2021; Urgoiti et al. 2022) and heterogeneity in shade tolerance (Morin et al. 2011; Zhang et al. 2012) are linked to the over-performance of mixed-species plots. We may expect to see likewise that variation in self-pruning strategies correlates with productivity.

Aside from a tree’s own traits, various characteristics of the surrounding tree community may be expected to influence the onset of self-pruning. The intensity of competition could alter the light threshold below which self-pruning occurs through correlative inhibition or alteration of the light compensation point. But even if a given tree species were always to initiate self-pruning at a fixed light threshold, the neighborhood could influence the height where that threshold is reached. A dense stand would be expected to have less light at any given height than a much sparser stand. Indeed, it is common for trees to have higher crown bases and lower crown depths in denser stands (Mäkelä 1997; Mäkelä & Vanninen 1998; Mäkinen 2002). Neighborhood characteristics could thus cause plasticity in the light threshold of self-pruning and in crown depth. Plastic adjustment in crown depth could conceivably either increase or decrease the extent of canopy packing in diverse tree mixtures relative to monocultures.

We posed questions and hypotheses about inter- and intraspecific variation in self-pruning among temperate trees within the context of a diversity experiment in southern Québec, Canada. Concerning interspecific variation, we posed the hypothesis that more shade-tolerant and resource-conservative species would initiate self-pruning at a deeper level of shade. Concerning intraspecific variation, we proposed that crowns that are well-lit at the top would also have a higher (i.e. less shaded) light threshold for self-pruning, in accordance with the notion of correlative inhibition. Consistent with this prediction, we also expected larger trees to have more light at the crown base (Nock et al. 2008). Lastly, we expected crown depth to decline with the intensity of competition in the local neighborhood.

We also posed hypotheses about the influence of variation in self-pruning on ecosystem function. We proposed that plots whose species vary in the light threshold of self-pruning would show greater productivity and canopy packing. We proposed further that intraspecific adjustment in crown depth from monocultures to mixtures would tend to increase canopy packing and strengthen the effect of diversity on canopy packing.

## Methods

### Site and experimental design

We conducted this study in the IDENT-Montréal experiment, which is part of the IDENT network of tree diversity experiments (Tobner et al. 2014) and the larger TreeDivNet (Paquette et al. 2018). The experiment was planted in spring 2009 on abandoned farmland in Ste-Anne-de-Bellevue, Québec, Canada (45° 25’ 30.1” N, 73° 56’ 19.9” W), west of the city of Montréal. Details about the site’s climate, soils, and preparation for experimental planting can be found in Tobner et al. (2016).

The experiment comprises 256 plots arrayed into four blocks. The plots were planted with varying species diversity and composition, drawing on a pool of 19 species. Here, we focus on plots which include only the 12 species native to North America—six evergreen conifers, one deciduous conifer, and five deciduous broadleaf species (Table 1). The distinct all-native species compositions included monocultures of all 12 species, 14 distinct two-species mixtures, 10 distinct four-species mixtures, and a 12-species mixture. Each composition was replicated once in each block, and position within the block was random.

**Table 1:** The twelve native species in IDENT-Montréal and their characteristics. The shade tolerance values come from Niinemets & Valladares (2006).

| Species | Species code | Family | Leaf habit | Leaf morphology | Shade tolerance |
| --- | --- | --- | --- | --- | --- |
| *Abies balsamea* (L.) Mill. | ABBA | Pinaceae | Evergreen | Needleleaf | 5.01 |
| *Acer rubrum* L. | ACRU | Sapindaceae | Deciduous | Broadleaf | 3.44 |
| *Acer saccharum* Marshall | ACSA | Sapindaceae | Deciduous | Broadleaf | 4.76 |
| *Betula alleghaniensis* Britton | BEAL | Betulaceae | Deciduous | Broadleaf | 3.17 |
| *Betula papyrifera* Marshall | BEPA | Betulaceae | Deciduous | Broadleaf | 1.54 |
| *Larix laricina* (Du Roi) K. Koch | LALA | Pinaceae | Deciduous | Needleleaf | 0.98 |
| *Picea glauca* (Moench) Voss | PIGL | Pinaceae | Evergreen | Needleleaf | 4.15 |
| *Picea rubens* Sarg. | PIRU | Pinaceae | Evergreen | Needleleaf | 4.39 |
| *Pinus resinosa* Aiton | PIRE | Pinaceae | Evergreen | Needleleaf | 1.89 |
| *Pinus strobus* L. | PIST | Pinaceae | Evergreen | Needleleaf | 3.21 |
| *Quercus rubra* L. | QURU | Fagaceae | Deciduous | Broadleaf | 2.75 |
| *Thuja occidentalis* L. | THOC | Cupressaceae | Evergreen | Scaleleaf | 3.45 |

Each plot was 4 × 4 m and had 64 trees planted in a 0.5 × 0.5 m grid. Within blocks, plots were separated by a 1.25 m buffer. Trees were planted at 1-2 years old. Tree species in a given plot were planted with equal frequency. By the time we conducted this study in 2018, the canopy had effectively closed in all plots.

### Self-pruning survey

We collected self-pruning measurements over one month in July and August 2018. We focused on two of the four blocks (A and D) and left out the 12-species plots. From each of the 72 selected plots, we randomly chose four living trees of each species when possible, excluding the plot edges. However, there were not always enough living trees to sample four per species per plot, particularly in four-species plots; this left us with *n* = 546 (out of a potential 640) living trees.

We made the key assumption that each tree’s crown base was defined by self-pruning of lower branches in the past, such that the present height and amount of light at the crown base could serve as measures of the threshold beneath which self-pruning occurs. We defined the crown base as the horizontal line above which there is relatively continuous live foliage that is typical for its species (USDA Forest Service 2011; Fig. 1). (This definition stands in contrast to those often used in fire ecology, where self-pruning is considered not to occur until dead branches physically detach from the trunk [Schwilk & Ackerly 2001].) On each tree, we measured the height of both the crown top and the crown base and estimated the crown radius as the average of four measurements in the cardinal directions from the central stem to the periphery. We calculated the crown depth as the vertical distance between the crown top and the crown base (Fig. 1A).

**Fig. 1:**
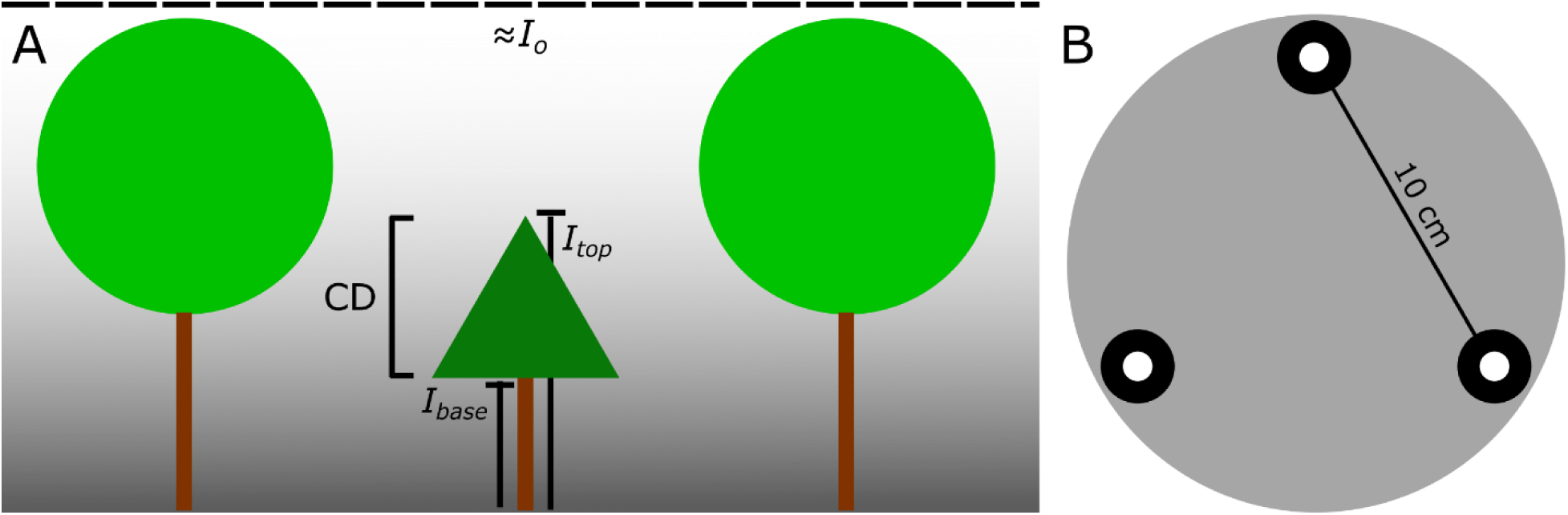
A diagram of our self-pruning measures. (A) For each focal tree, we calculated the crown depth (CD) as the difference between the height at the top and base of the crown. We used an apparatus depicted in (B), represented here by the T-shaped figures, to measure the amount of light at the base and top of the crown (*I_base_* and *I_top_*). These values were compared to the amount of light in an adjacent open environment (*I_o_*), a proxy for light above the canopy of the entire experiment (dashed line). (B) To measure light, we used three light sensors fixed to a plate in an equilateral triangle. This plate was mounted on an extensible pole, which held it parallel to the ground.

We also characterized the general availability of light (as photosynthetic photon flux density) at both the crown top and the crown base. Following Messier & Puttonen (1995), we calculated the ratio between the amount of light at each site of interest and in an adjacent open environment, which we took as representative of conditions above the canopy. All measurements were conducted under clear-sky conditions due to the rarity of consistently overcast conditions; this method can estimate average light availability with little bias, but with greater temporal and spatial noise compared to overcast conditions (Messier & Puttonen 1995). We only collected measurements from 9:00-16:00 to avoid times when the sun was close to the horizon (median solar zenith angle: 36.6°; 2.5^th^-97.5^th^ percentile: 25.5°-55.2°).

We measured a time series of light in the open (*I_o_*) using an LI-190R sensor mounted on a tripod and connected to a data logger. We measured light using three LI-190R sensors (LI-COR Biosciences, Lincoln, NE, USA) arrayed in an equilateral triangle with a side length of 10 cm on a round wooden plate (Fig. 1) mounted on an extensible pole. For each selected tree, we went in each of the four cardinal directions from the trunk, stopping short of the edge of the crown base. Just below the base in each direction, we took simultaneous readings from the three sensors three times, rotating the plate between readings, to measure *I_base_*. Each measurement was thus an average of 36 individual sensor readings (4 directions × 3 rotations × 3 sensors). We then repeated this procedure just above the crown top to measure *I_top_*, although for some trees that were visibly the tallest within a large neighborhood, we assumed *I_top_* to be equal to *I_o_*. We summarized the properties that determine light at the crown base and top by calculating the values *L_base_ =* ln(*I_base_*/*I_o_*) and *L_top_* = ln(*I_top_*/*I_o_*), extracting *I_o_* from the time-series based on measurement time. These log-light fractions reach a maximum of 0 under open conditions. The formulas are based on an idealization of light transmission through the canopy as an exponential decay described by the Beer-Lambert law (Ponce de León & Bailey 2019). Besides this theoretical justification, the log-transformation also helps to linearize the relationships between these quantities and their predictors.

Both *L_base_* and crown depth can serve as measures of self-pruning, but with different functions and interpretations. *L_base_* describes the threshold of light needed to cause self-pruning, but the height above the ground where this threshold is reached depends on the pattern of light extinction through the canopy, which could vary strongly across neighborhoods even if *L_base_* were constant. Crown depth integrates potential variation in both *L_base_* and the light environment, which together shape the geometry of the crown.

### Growth survey

Every autumn from the experiment’s inception, we coordinated a survey to measure the basal diameter (15 cm above the ground) of every live tree in the experiment. Based on the 2018 survey, we estimated productivity for each native-species plot, including 12-species plots and all four blocks, using the basal area of the inner 6 × 6 trees (excluding edges). Trees that died by 2018 were considered to make no contribution to basal area.

We also characterized the competitive environment of each focal tree’s neighborhood using the neighborhood competition index, calculated as *NCI* = Σi *Ai* / *Di*, where for each tree *i* in the neighborhood, *A_i_* is the stem cross-sectional area and *D_i_* is the distance. In effect, this index simplifies the more complex NCI of Canham et al. (2004) by assuming that a tree’s competitive effect is independent of its species identity, and by setting the parameters *α* and *β*—exponents of tree diameter in the numerator and distance in the denominator—to 2 and 1. We considered these simplifications appropriate since our aim is inference rather than prediction. We defined the neighborhood to include all trees within a radius of 1.1 m from the focal tree (the 90^th^ percentile of crown radius measurements).

### Functional identity and diversity

To describe the functional identity of each species, we used a set of five traits describing plant economics: specific leaf area (SLA), leaf nitrogen concentration (%N), leaf dry matter content (LDMC), wood density (WD), and specific root length (SRL). These traits had previously been compiled from trait databases for IDENT-Montréal species (Belluau 2020). We used principal components analysis on the correlation matrix to identify major axes of species-level variation (Fig. S1). The first principal component explained 56.5% of trait variation, so we took species’ positions along this axis to represent their functional identity. The axis separated the evergreen needleleaf species at low values from the deciduous broadleaf species at high values, with the one deciduous needleleaf species (*L. laricina*) in between (Urgoiti et al. 2022); higher values correspond to higher SLA, %N, WD, and SRL. We refer to this axis as ‘acquisitiveness’ based on correspondence to the global whole-plant economic spectrum (Díaz et al. 2016). While the loading for wood density is contrary to expectations based on global patterns (Díaz et al. 2016), it is consistent with a typical split between coniferous softwoods and broadleaf hardwoods in North American temperate forests. Past studies in the experiment noted that deciduous species had greater aboveground growth than evergreen species, as evaluated from the annual survey (Tobner et al. 2016; Urgoiti et al. 2023b). Because this division persisted at the time of our measurements, acquisitiveness was to some degree confounded with tree size across species. For each focal tree in our self-pruning survey, we described the functional identity of trees within the 1.1 m radius neighborhood by calculating the community-weighted mean (CWM) of acquisitiveness (‘neighborhood acquisitiveness’), with weights determined by the number of trees planted of each species.

Aside from the trait-based dimension of acquisitiveness, we also described focal tree strategies using the shade tolerance index of Niinemets & Valladares (2006), which was compiled on a 1-5 scale based on empirical measurements and expert opinion. Likewise, we also used previously compiled leaf lifespan for each species (Belluau 2020). Although leaf lifespan is often considered a leaf economic trait (Wright et al. 2004), we considered it separately here to investigate relationships between leaf and branch longevity.

While we carried out analyses of inter- and intraspecific variation in self-pruning at the scale of individuals and their neighborhoods, we examined their influence on ecosystem function at the scale of entire plots. To enable these analyses, we calculated various metrics of diversity at the plot-level, including the diversity of functional traits, of shade tolerance values, and of *L_base_*. We calculated diversity of *L_base_* using species means across all plots to make it analogous to the functional traits and shade tolerance values, which were also available only at the species level. In these calculations, we used the ^q^D(TM) diversity metric proposed by Scheiner et al. (2017), with weights determined by the number of trees of each species planted among the plot’s inner 6 × 6 trees. This metric corresponds to the effective number of species with maximal distinctness. The Hill number *q* was set to 1, resulting in measures analogous to the exponential of Shannon diversity (Jost 2006). To calculate functional trait ^q^D(TM), we used a matrix of Euclidean distances among species, calculated with the five aforementioned functional traits (*z*-standardized).

### Statistical analyses

We conducted all analyses in *R* v. 4.4.0 (R Core Team 2024). To examine how self-pruning behavior is related to other aspects of trees’ strategies, we used ordinary least-squares (OLS) regression to evaluate relationships between species-level properties (acquisitiveness, shade tolerance, and leaf lifespan) and species mean *L_base_*.

To understand drivers of variation among individual trees, we tested the influence of four variables on *L_base_*. Two of these variables were properties of the focal individual itself (height and light fraction at the crown top) and two were properties of its neighborhood (NCI and CWM acquisitiveness). Using package *lme4* v. 1.1.35.3 (Bates et al. 2015), we built mixed-effects models testing how light fraction at the crown base is influenced by the interactive effects between one individual-level variable at a time and species identity, with plot as a random intercept. We used *lmerTest* v. 3.1.3 (Kuznetsova et al. 2017) to calculate *p*-values using Satterthwaite’s method to estimate degrees of freedom, noting that such estimates should be interpreted conservatively. We used analysis of covariance (ANCOVA) with Type III sums of squares to test the significance of each main effect and their interaction. Under the hypothesis of correlative inhibition, we expected *L_top_* to have a positive effect on *L_base_*. Whether or not there was a significant interaction between species and each individual-level variable, we extracted the species-specific slopes using package *emmeans* v. 1.10.3 (Lenth 2024) and used OLS regression to evaluate whether these slopes covaried with shade tolerance or acquisitiveness.

We likewise tested the influence of these same four variables on crown depth using ANCOVAs with identical structure to the models of *L_base_*. Crown depth was strongly dependent on tree height, which varied considerably among species, making it challenging to interpret species-specific slopes. For this reason, we were primarily interested in the significance and direction of the main effects of the four individual variables rather than their interactions with species identity. Accordingly, we also did not analyze trends in species-specific slopes.

We were interested in whether diversity in *L_base_*—considered as a putative functional trait at the species level—could explain productivity across plots, analogous to previous studies using better-studied functional traits (Bongers et al. 2021; Urgoiti et al. 2022). We used OLS regression to test whether diversity in functional traits (not including *L_base_* or leaf lifespan), shade tolerance, and *L_base_* were correlated with productivity at the plot level. In this analysis alone, we used all native-species plots (including those with twelve species) in all four experimental blocks, since the analysis depended only on species means rather than individual measurements of *L_base_*.

### Simulations of canopy packing

In evaluating the influence of intraspecific variation in self-pruning on canopy packing, we used crown depth as a metric of how neighborhoods influenced crown shape via self-pruning. We relied on a simple, stylized representation of crown structure developed by Purves et al. (2007) and adopted by Jucker et al. (2015) in the context of canopy packing. In this representation, tree crowns are defined by the positions of their base and top, their radius, and a shape parameter that influences concavity. This representation makes it tractable to estimate the volume of individual crowns, or even the hypothetical overlap between two crowns (as in Williams et al. [2017]). We fixed the shape parameter at the species level using estimates provided by Purves et al. (2007). Further details on calculating crown volume are found in the Supporting Information.

To evaluate how self-pruning alters canopy packing, we simulated crown volume under two scenarios: (1) one where trees in each plot show the same crown depth as in the monoculture of their species, and (2) one where trees may plastically adjust their crown depth when growing in mixtures. To carry out scenario (1), we simulated 500 crowns of each species in each plot; the radius was sampled from among the trees of that species in that plot measured in the self-pruning survey, but the depth was sampled from monocultures of the same species in the same block. To carry out scenario (2), we sampled both the crown depth and radius independently from the focal plot. (When the focal plot was a monoculture, the two scenarios were of course identical.) We took the second scenario as an estimate of the actual crown volume in the plot (Fig. S2). We performed this procedure for each plot included in the self-pruning survey. Further details on these simulations are presented in the Supporting Information.

For both scenarios, we scaled up individual crowns to estimate the total crown volume per unit area in each plot’s inner 6 × 6 trees (9 m^2^). Since our self-pruning survey only included living trees, we used the growth survey data to calculate the fraction of living trees of each species in each plot in 2018. We then estimated the total crown volume per unit area (*V*, in m^3^ m^-2^ or simply m) in each plot as:

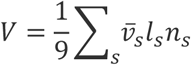

where ν̅_s_ is the mean crown volume of measured (living) trees of species *s* in the plot, *l_s_* is the fraction of trees of species *s* in the plot that were still alive, and *n_s_* is the number that were planted at the outset of the experiment.

## Results

### Species-level predictors of self-pruning

Among the individual-level variation in the log-light fraction at the crown base (*L_base_*), 51.3% could be explained by species identity alone. The *L_base_* values were lower among the six evergreen species than the six deciduous species. Given that evergreen species were also less acquisitive than deciduous species, there was a strong positive species-level correlation between *L_base_* and acquisitiveness (*t*(10) = 4.36, *R^2^*= 0.655, *p* = 0.001; Fig. 2). Shade tolerance was not linked to leaf habit, but showed a moderate negative correlation with *L_base_* (*t*(10) = -2.64, *R^2^* = 0.411, *p* = 0.025). Shade tolerance and acquisitiveness were uncorrelated (*t*(10) = -0.63, *R^2^* = 0.038, *p* = 0.542), and the two variables could additively explain by far most of the variation in *L_base_* (*R^2^* = 0.897). In addition, *L_base_* had a strong negative correlation with log-leaf lifespan both across all species (*t*(10) = -7.40, *R^2^* = 0.846, *p* < 0.001) and among the evergreen species (*t*(4) = -4.32, *R^2^* = 0.824, *p* < 0.001; Fig. 2).

**Fig. 2:**
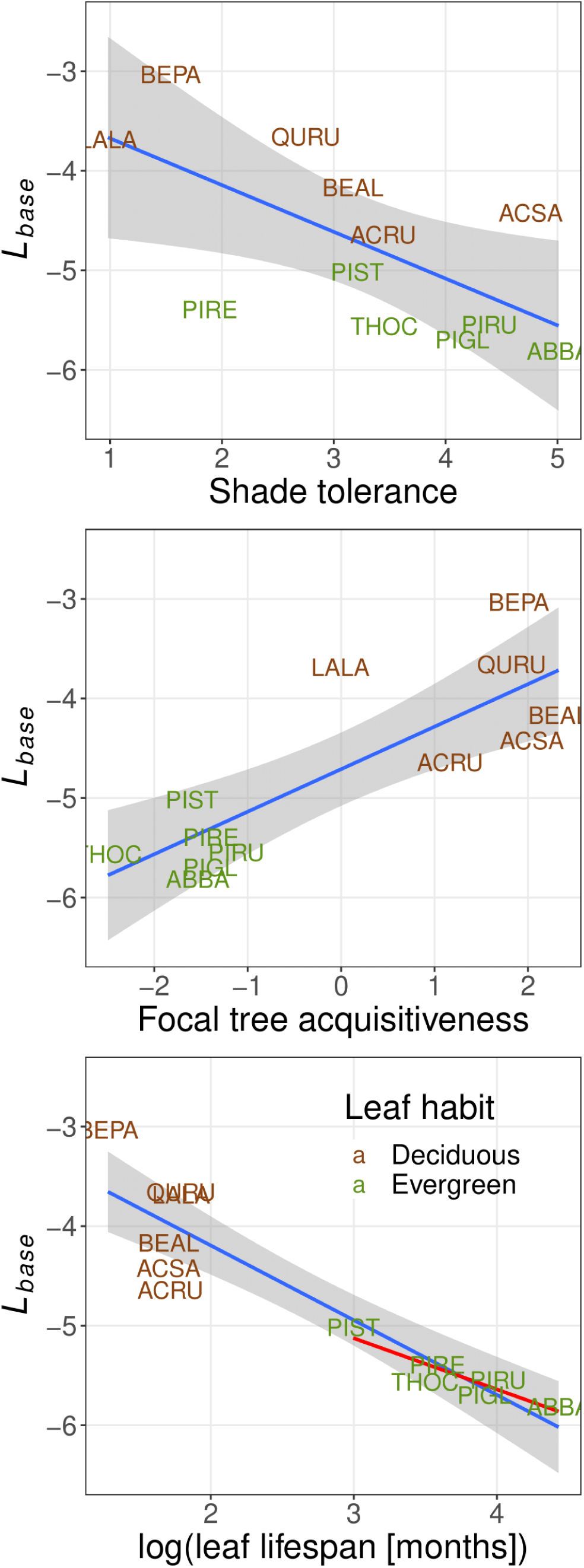
Species-level correlations between the light threshold of self-pruning (*L_base_*) and shade tolerance (top), acquisitiveness (middle), and leaf lifespan (bottom). Labels are species codes, which are found in Table 1. Blue lines represent OLS regression lines, with a 95% confidence interval shown in gray. The red line in the bottom panel is the OLS regression line for only evergreen species, with no confidence interval shown.

### Neighborhood-level predictors of self-pruning

We considered the interactive effects of species identity and each of four individual-level predictor variables (neighbor acquisitiveness, NCI, focal tree height, and *L_top_*) on *L_base_* in our mixed-model ANCOVA analyses. The four variables had clear but noisy correlations with each other within species (Fig. S3), most likely because more acquisitive neighbors increased competition, suppressed the growth of the focal individual, and reduced *L_top_*. There was strong evidence for a species × height interaction and weaker evidence for a main effect of height (Table 2). In the species × *L_top_* model there were strong main effects of species and *L_top_*, but much more modest evidence for an interaction. In the species × neighbor acquisitiveness model, we found evidence for a main effect of species and modest evidence for an interaction with acquisitiveness, but no main effect of acquisitiveness. In the species × NCI model, there was only a significant effect of species.

**Table 2:** Output from analysis of covariance (ANCOVA) models with Type III sums of squares designed to test how log-transformed light fraction at the crown base (*L_base_*) and crown depth (CD) are influenced by each of four individual/neighborhood-level variables (continuous), focal species identity (categorical), and their interactions. All models included a random effect for experimental plot, which is left out from the model structure for concision. As is typical for mixed-effects models, denominator degrees of freedom (df) varied for terms within a model and were estimated using Satterthwaite’s method. Abbreviations: neighbor acq = neighbor acquisitiveness; NCI = neighborhood competition index; *L_top_* = log-transformed light fraction at the crown top.

| Model structure | Term | Sum of squares | Mean squares | Numerator df | Denominator df | *F* | *p*-value |
| --- | --- | --- | --- | --- | --- | --- | --- |
| $L_{base} \sim$<br>neighbor acq<br>* species | neighbor acq | 0.249 | 0.249 | 1 | 414.7 | 0.425 | 0.515 |
| | species | 99.9 | 9.084 | 11 | 460.2 | 15.510 | $< 10^{-3}$ |
|  | interaction | 11.6 | 1.053 | 11 | 467.3 | 1.798 | 0.052 |
| $L_{base} \sim$ NCI *<br>species | NCI | 0.394 | 0.394 | 1 | 489.7 | 0.655 | 0.419 |
| | species | 60.3 | 5.48 | 11 | 476.4 | 9.114 | $< 10^{-3}$ |
|  | interaction | 3.07 | 0.280 | 11 | 508.4 | 0.465 | 0.925 |
| $L_{base} \sim$ tree<br>height *<br>species | tree height | 2.89 | 2.89 | 1 | 455.3 | 4.906 | 0.027 |
|  | species | 4.83 | 0.439 | 11 | 514.8 | 0.744 | 0.696 |
| | interaction | 22.4 | 2.04 | 11 | 495.5 | 3.454 | $< 10^{-3}$ |
| $L_{base} \sim L_{top}$ *<br>species | $L_{top}$ | 5.48 | 5.48 | 1 | 518.2 | 9.653 | 0.002 |
| | species | 68.0 | 6.18 | 11 | 475.6 | 10.890 | $< 10^{-3}$ |
|  | interaction | 9.91 | 0.901 | 11 | 512.0 | 1.587 | 0.099 |
| CD ~<br>neighbor_acq<br>* species | neighbor acq | 70948 | 70948 | 1 | 388.7 | 8.884 | 0.003 |
| | species | 1190804 | 108255 | 11 | 444.1 | 13.555 | $< 10^{-3}$ |
|  | interaction | 136914 | 12447 | 11 | 452.6 | 1.559 | 0.108 |
| CD ~ NCI *<br>species | NCI | 47536 | 47536 | 1 | 460.6 | 5.784 | 0.017 |
| | species | 464877 | 42262 | 11 | 461.0 | 5.142 | $< 10^{-3}$ |
|  | interaction | 60639 | 5513 | 11 | 510.2 | 0.671 | 0.767 |
| CD ~ tree<br>height *<br>species | tree height | 1608601 | 1608601 | 1 | 487.3 | 608.872 | $< 10^{-3}$ |
| | species | 166282 | 15117 | 11 | 512.7 | 5.722 | $< 10^{-3}$ |
|  | interaction | 58917 | 5356 | 11 | 506.5 | 2.027 | 0.024 |
| CD ~ $L_{top}$ *<br>species | $L_{top}$ | 1199701 | 1199701 | 1 | 488.3 | 225.623 | $< 10^{-3}$ |
| | species | 446413 | 40583 | 11 | 424.2 | 7.632 | $< 10^{-3}$ |
| | interaction | 211907 | 19264 | 11 | 507.6 | 3.623 | $< 10^{-3}$ |

Even though we only found any evidence for species variation in slopes for two individual-level predictor variables, we extracted the species-specific slopes from the mixed models to evaluate whether there were any suggestive patterns for further exploration. There was no evidence for any correlations between these slopes and shade tolerance (all *p* > 0.05); however, there were varying degrees of evidence for correlations with the focal species’ acquisitiveness (Fig. 3). There was no significant correlation for *L_top_* (*t*(10) = 1.59, *R^2^* = 0.202, *p* = 0.143), but progressively stronger correlations for NCI (*t*(10) = -3.298, *R^2^* = 0.521, *p* = 0.008), neighbor acquisitiveness (*t*(10) = -3.759, *R^2^*= 0.586, *p* = 0.004), and tree height (*t*(10) = 6.058, *R^2^* = 0.786, *p* < 0.001). In particular, the most acquisitive species had a greater *L_base_* when their neighborhood had lower NCI and acquisitiveness, and when they themselves had greater height and light at the crown top as a result. This trend was reversed among the most conservative species, which had a lower *L_base_* when their neighborhood had lower NCI and acquisitiveness and when they had greater height. However, no species had highly negative correlations between *L_top_* and *L_base_*.

**Fig. 3:**
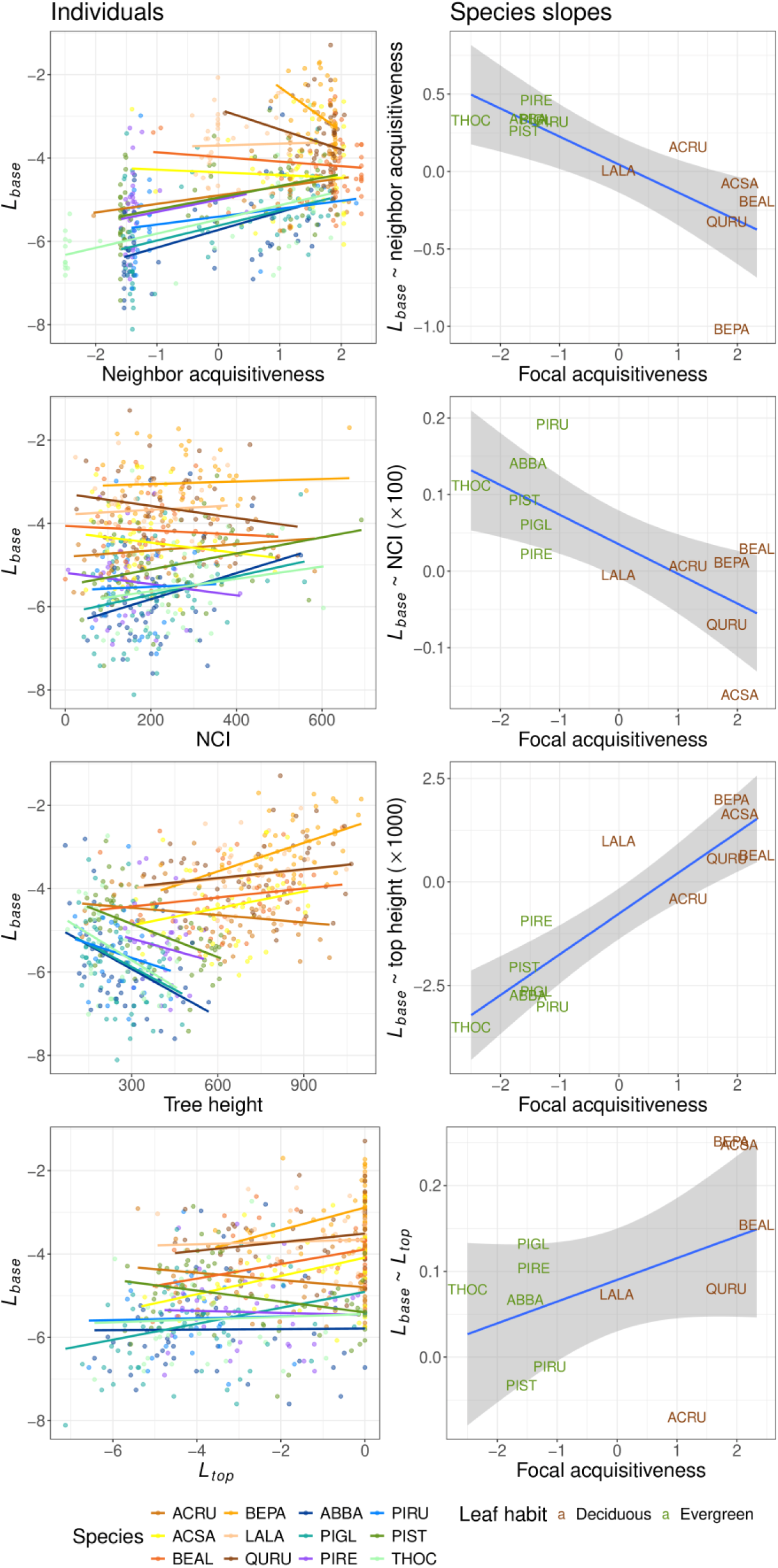
Relationships between individual-level variables and *L_base_* across species. The individual-level variables are neighbor acquisitiveness (top row), the neighborhood competition index (NCI; second), the focal tree’s height (third), and the light availability at the crown top (*L_top_*; bottom). Left panels show the individual-level relationships with species-specific best-fit lines, while right panels show species’ slopes in relation to their acquisitiveness. The slopes of the lines in the left panels are not exactly identical to the slopes plotted along the *y*-axis in the right panels, since the former are OLS regression slopes and the latter are based on mixed-effects models that also include a random intercept for plot.

There was strong evidence that crown depth was greater for individuals with lower NCI, lower neighbor acquisitiveness, higher focal tree height, and higher light at the crown top (Table 2; Fig. 4). Besides these main effects, species identity and (for all models except the one with NCI) the interaction term also had significant effects, implying that species varied in crown depth even accounting for their other individual and neighborhood characteristics.

**Fig. 4:**
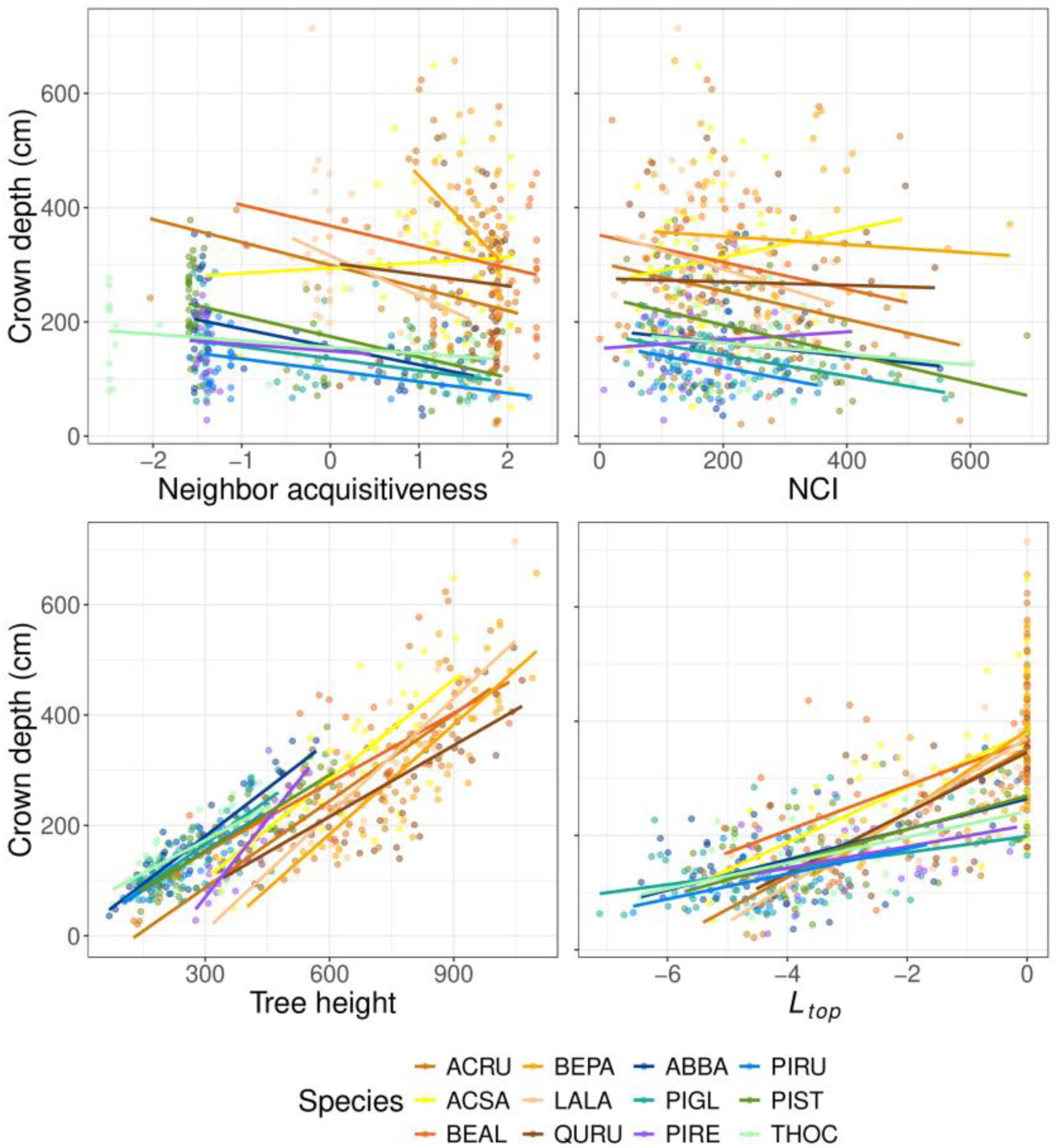
Relationships between individual-level variables and crown depth across species. The individual-level variables are neighbor acquisitiveness (top), the neighborhood competition index (NCI; second), the focal tree’s height (third), and the light availability at the crown top (*L_top_*; bottom). The best-fit lines are derived from species-specific OLS regressions.

### Self-pruning and productivity

We used plot-level diversity (*^q^*D(TM)) in functional traits, shade tolerance, and *L_base_* as predictors of plot basal area. Log-transforming our diversity measures nearly always improved model fit. After this log-transformation, diversity in *L_base_* was better correlated with basal area (*t*(146) = 5.71, *R^2^* = 0.182, *p* < 0.001; Fig. 5) than diversity in functional traits (*t*(146) = 3.81, *R^2^* = 0.090, *p* < 0.001) or diversity in shade tolerance (*t*(146) = 3.42, *R^2^* = 0.074, *p* < 0.001). Similar results hold when considering the net biodiversity effect (NBE) on productivity, defined as the overperformance of mixtures relative to monoculture-based expectations (Fig. S4; see Supporting Information for further details).

**Fig. 5:**
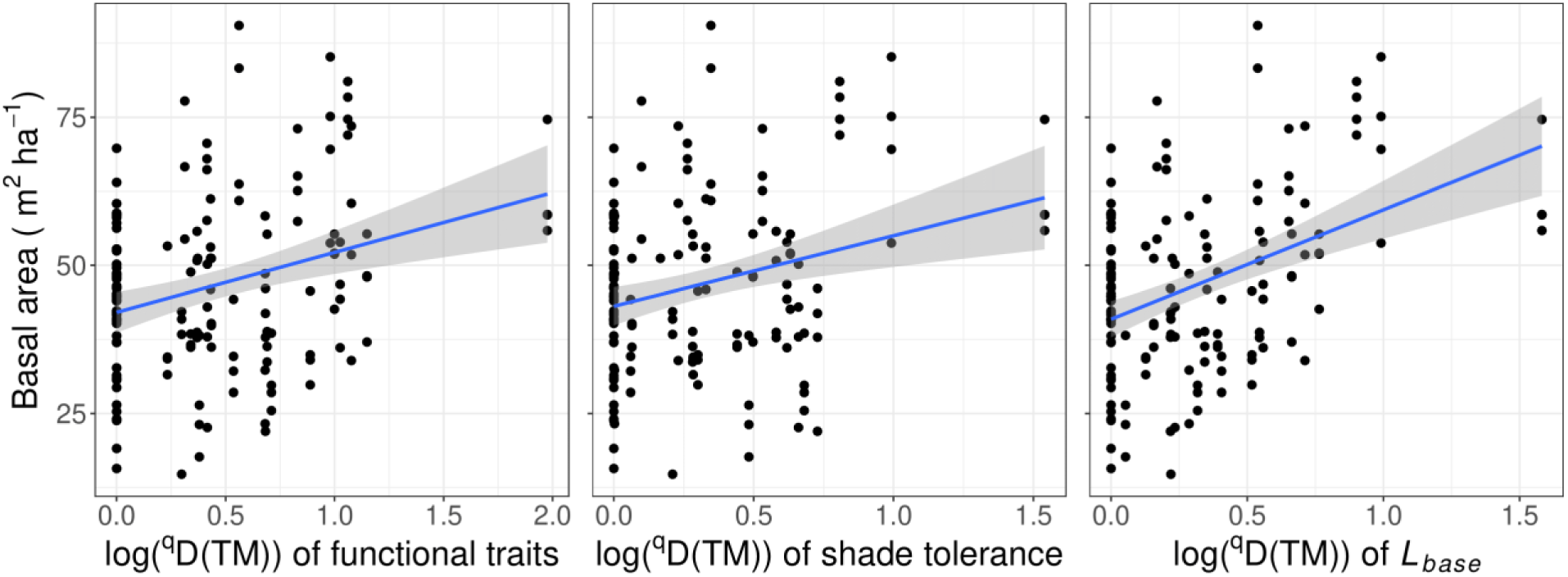
Correlations between basal area and diversity of functional traits (left), shade tolerance (middle), and *L_base_* (right) among plots. Blue lines are OLS regression lines, with 95% confidence intervals shaded in gray.

### Canopy packing

Estimated total crown volume per plot (based on simulations under the scenario with plasticity) was moderately correlated with basal area (*t*(67) = 5.35, *R^2^* = 0.299, *p* < 0.001). Crown volume increased modestly with the diversity (log-transformed ^q^D(TM)) of *L_base_* (*t*(67) = 4.45, *R^2^* = 0.228, *p* < 0.001; Fig. 6) and of functional traits (*t*(67) = 3.81, *R^2^* = 0.178, *p* < 0.001).

**Fig. 6:**
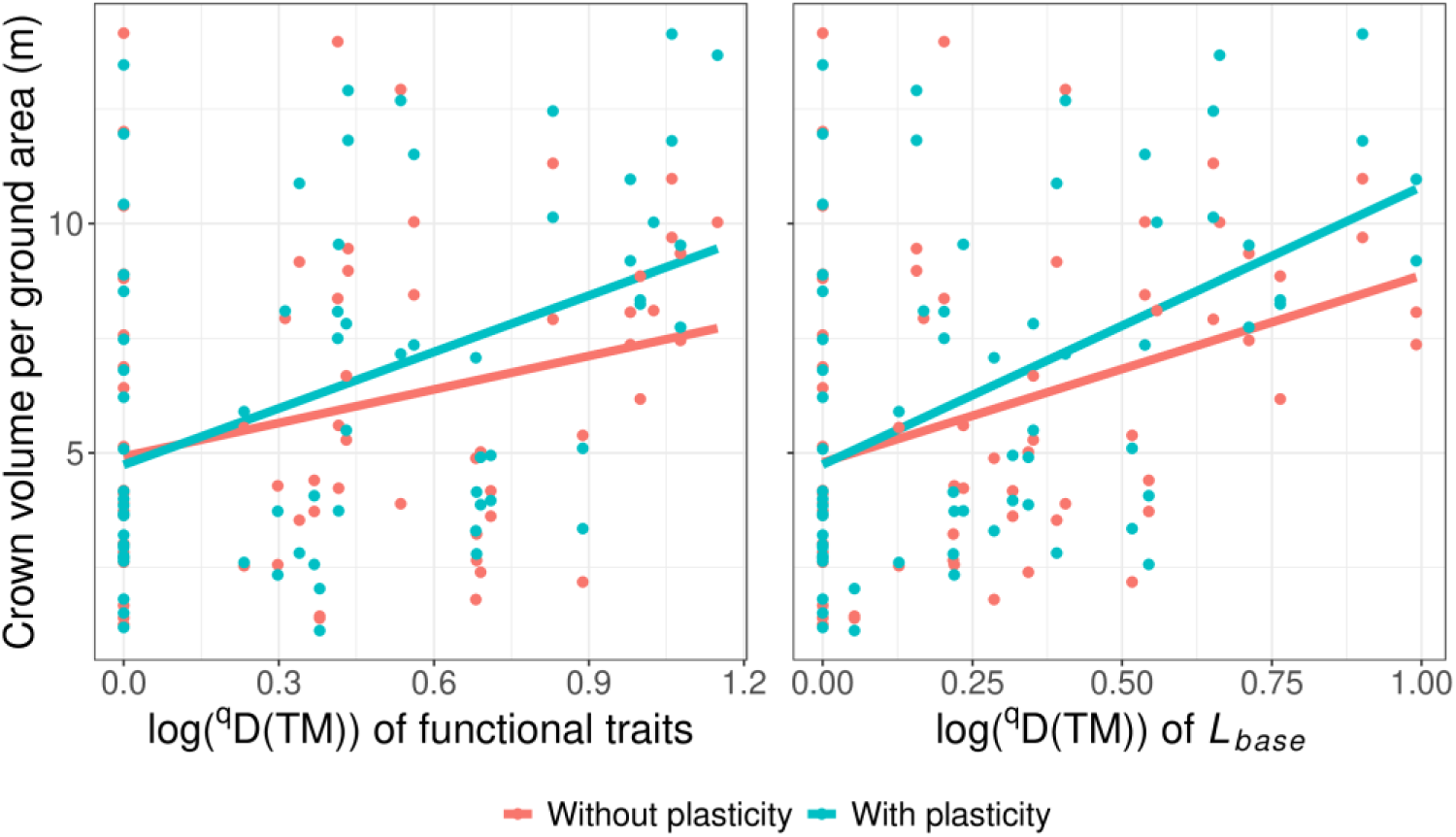
The relationship between total crown volume and diversity in functional traits (left) or *L_base_* (right) in simulations where plasticity in crown depth between monoculture and mixtures is or is not allowed. Lines represent OLS regression fits.

To evaluate whether the plastic adjustment of crown depth influences canopy packing, we compared simulated mixture plots where crown depths were drawn from monoculture plots (without plastic adjustment) to those where crown depths were drawn from the focal plot (with plastic adjustment). Simulated mixture plots without plastic adjustment showed less total crown volume than those with plastic adjustment (paired *t*-test; *t*(44) = 3.05, *p* = 0.004, mean difference = 0.810 m). The greatest source of this disparity was in the most productive mixtures, which yielded as much or more crown volume than the most productive monocultures in actual data, but considerably less in simulations (Fig. 6). As a result, the relationship between total canopy packing and diversity in functional traits was attenuated in these simulations without plastic adjustment in crown depth (*t*(67) = 2.35, *R^2^* = 0.076, *p* = 0.022) or *L_base_* (*t*(67) = 3.10, *R^2^* = 0.125, *p* = 0.003). Similar results hold when considering the net biodiversity effect on canopy packing (Fig. S5; see Supporting Information for further details).

## Discussion

We found considerable variation in the threshold of light availability below which trees pruned their lower branches. This value, which we called *L_base_*, was lower for more conservative and shade-tolerant species, and plots whose species had greater diversity in *L_base_* were more productive. However, there were also notable intraspecific patterns: while acquisitive species had lower *L_base_* with acquisitive neighbors, conservative species had lower *L_base_* with conservative neighbors. Moreover, there was a general tendency for *L_base_* to be higher in crowns with sunlit tops, consistent with predictions based on correlative inhibition. Most species had deeper crowns when surrounded by smaller and more conservative neighbors. In simulations, we found that this plasticity in crown depth increased the total amount of crown volume in diverse plots, strengthening the relationship between crown volume and functional diversity. Our findings suggest that self-pruning plays a role in defining trees’ functional strategies, and that diversity in self-pruning strategies may contribute to biodiversity-ecosystem functioning relationships in light-limited mesic forests.

### Interspecific variation in self-pruning

A large body of research has emphasized the role of shade tolerance in either self-pruning (Schoonmaker et al. 2014) or related aspects of crown and stand structure, including crown shape (Poorter et al. 2009), canopy light transmission (Canham et al. 1994; Valladares & Niinemets 2008), leaf area index (Niinemets 2010), and delays in self-thinning (Urgoiti et al. 2023). We found that *L_base_* did correlate with shade tolerance across species, but it correlated even more strongly with our index of resource-acquisitiveness. The directions of these relationships agreed with our predictions, which were based on the assumption that branches die when they become carbon-depleted. Furthermore, although shade tolerance is often taken to be strongly influenced by economic traits like dark respiration (Baltzer & Thomas 2007a, b; Lusk & Jorgensen 2013), shade tolerance and acquisitiveness were entirely uncorrelated among our species.

Future comparative work on the drivers of self-pruning may benefit from directly measuring the light compensation point and dark respiration, rather than taking them to be proxied by broad axes of plant functional variation. In our experiment, acquisitiveness was tightly linked with both leaf habit/type and focal tree size, since the most acquisitive species were deciduous and fast-growing. These links raise the question of whether acquisitive species only have higher *L_base_* due to their size rather than their traits. Were this the case, the largest and least-shaded trees among the conservative evergreen species would approach the *L_base_* of the acquisitive deciduous species; by contrast, actual patterns of plasticity suggest that these groups diverge with increasing tree size. This potential for confounding could be addressed more directly in the future by measuring isolated trees of similar sizes.

Leaf lifespan was also tightly linked to average *L_base_* across species. There may be something intuitively logical about this link, given that *L_base_* could exert a strong influence on branch longevity; for example, holding constant a stand’s timeline of canopy closure and light extinction, a species with lower *L_base_* would be expected to retain its lower branches for a longer period. In a deeper sense, both leaf lifespan and *L_base_* may be underpinned by economic traits that determine how long it takes for the leaf and branch to fall below their respective light compensation points (Givnish et al. 1988; Reich et al. 2009).

Recent work has aimed to identify crown economic traits that define fundamental axes of variation in tree architecture, analogous to the leaf economic spectrum (McNeil et al. 2023). We propose that *L_base_* could serve as a simple crown economic trait to describe trees’ strategies in light-limited environments. Indeed, it may serve as both a response and effect trait (*sensu* Lavorel & Garnier 2002): a response trait in that it has a role in determining how much leaf area a tree retains given the light environment created by its neighbors, and an effect trait in that it determines how much shade the tree casts on any plants below itself. Species with lower *L_base_* may be able to persist under more closed canopies, and when dominant may be more likely to prevent establishment of shade-intolerant species in the understory (Rees et al. 2001; Craine & Reich 2005), helping to drive the ratchet of succession. This interpretation depends in part on whether species rank similarly in *L_base_* across environments. We found that there are strong and fairly consistent interspecific patterns in *L_base_* despite plastic variation, which suggests that the trait could help explain species turnover during succession in light-limited forests.

Our results leave open the question of whether *L_base_* is redundant with known dimensions of variation in plant function, including shade tolerance and the plant economic spectrum. If it is, these economic traits may be more strongly intertwined with canopy properties than currently recognized. Regardless, it is unlikely to simply represent a general deciduous-evergreen split; for example, some very shade-intolerant pines transmit large fractions of light (>25%) even at fairly high stocking densities (Battaglia et al. 2003; Knapp et al. 2016). A broader sampling of species could help resolve *L_base_*’s links to better-studied traits.

### Plasticity in self-pruning

Besides the clear trends across species, we also found patterns of plasticity in *L_base_* and crown depth across neighborhoods. We found that *L_base_* showed contrary relationships with neighborhood characteristics among acquisitive and conservative species. For acquisitive species, *L_base_* became more negative with more acquisitive and larger neighbors. Since large neighbors reduced focal tree height—presumably via competition—*L_base_* was also more negative among smaller focal trees. For conservative species, *L_base_* was less negative with more acquisitive and larger neighbors, and thus less negative among smaller focal trees.

The rise in *L_base_* with size among more acquisitive species is consistent with findings that leaf area index declines and light transmission increases as deciduous tree crowns grow (Nock et al. 2008).

Likewise, shade tolerance tends to decline with ontogeny (Kneeshaw et al. 2006). These patterns may be explained by increasing respiratory load (Givnish et al. 1988; Sendall et al. 2015) or by correlative inhibition manifesting in increased allocation towards horizontal growth in sunlit parts of the canopy (Nock et al. 2008), among other possibilities. It is harder to plausibly explain why *L_base_* becomes more negative with size among more conservative species. Because these species tend to be much less dominant in mixture plots, they may experience greater shading from acquisitive neighbors alongside self-shading—especially the smallest individuals with the least negative *L_base_*. Some of these small individuals of conservative species have crown bases nearly at ground level (Fig. S6), so they may not have initiated self-pruning at all.

Our results show modest evidence that trees had more light at the crown base when they had more light at the crown top. This pattern is consistent with correlative inhibition, in which trees whose crowns are at least partly sunlit tend to drop their lower branches at a brighter light threshold than trees that are fully shaded. However, there may be other conceivable explanations for this trend: for example, if one of a focal tree’s neighbors suddenly dies, it would increase the amount of light at both the crown top and the crown base, but not due to self-pruning strategies. Although Schoonmaker et al. (2014) had proposed that correlative inhibition may be strongest in shade-intolerant species because their strategies to acquire and preempt light require stronger preferential allocation to well-lit parts of the canopy, we did not find evidence for this effect. More direct manipulations may be needed to examine competing explanations and decisively test how the strength of correlative inhibition varies among species (Henriksson 2001; Schoonmaker et al. 2014).

All species showed declines in crown depth as the neighborhood environment became more competitive. This result would be expected even in the absence of plasticity in *L_base_*, since (1) competitive environments reduce the focal tree’s top height, and (2) more competitive environments result in greater light extinction, which would cause the focal tree’s *L_base_* to be reached at a greater height. However, the plasticity in *L_base_* can be assumed to contribute to variation in crown depth; if they had had constant *L_base_*, the most acquisitive species would presumably have had steeper declines in crown depth with increasing neighbor size or acquisitiveness, while conservative species would have had shallower declines.

### Self-pruning, diversity effects, and canopy packing

Past research in IDENT-Montréal has shown that species mixtures show a positive average net biodiversity effect (NBE)—they grow more than expected based on monocultures of their constituent species. In early years, this effect was weak and due almost entirely to the over-performance of high-yielding broadleaf species, which benefited from competitive release in mixtures with less productive species (Tobner et al. 2016). Across plots, the magnitude of the NBE was closely related to the increase in crown complementarity from monocultures to mixtures (Williams et al. 2017), and diverse plots had greater light interception (Rissanen et al. 2019). However, by the time of the self-pruning survey in 2018, there were more consistently positive NBEs that were driven by the over-performance of both high- and low-yielding species, and the NBE was largest in plots with high diversity in plant economic traits (Urgoiti et al. 2022). There was also strong evidence for transgressive overyielding, in which certain mixtures are more productive than the most productive monocultures.

We found that diversity in *L_base_* (as a measure of self-pruning strategies) was better correlated with plot basal area and NBEs than were diversity in functional traits or shade tolerance (Fig. 5; Fig. S4). Likewise, diversity in functional traits and especially in *L_base_* were positively related to canopy packing (Fig. 6; Fig. S5). Moreover, our simulations showed that the relationship between canopy packing and diversity was strengthened by plasticity in crown depth. The most acquisitive and productive species in the experiment had much deeper crowns in mixtures with less acquisitive species than in the highly competitive environment of monocultures (Fig. 4). As a result, mixtures with high diversity in functional traits or *L_base_* produced about as much crown volume as the most productive monocultures, even though in the absence of plasticity in crown depth they would have produced much less. However, plasticity in crown depth is not solely responsible for the positive relationship between diversity and canopy packing, since the relationship appeared even in simulations without plasticity.

In general, the influence of diversity on productivity likely arises from multiple mechanisms that influence species net interactions. However, in a dense, closed-canopy forest undergoing self-thinning (Urgoiti et al. 2023b), competition for light can generally be taken as one of the dominant mechanisms. Under competition for light, the influence of species interactions on total productivity arises from a mixture of two effects: (1) competitive relaxation, in which light niche partitioning allows the community to achieve greater total light interception; and (2) competitive imbalance, in which the stark asymmetry of light competition causes most of the light to be intercepted by the most dominant species (Yachi & Loreau 2007). Competitive imbalance usually has a negative influence, in that the stunted growth or death of deeply shaded species results in a loss of productivity. However, given that our results and past findings at IDENT-Montréal show evidence for transgressive overyielding and for over-performance of less-dominant species (Fig. 3; Urgoiti et al. 2022), it appears that competitive imbalance may not have been strong enough to compromise total growth.

In IDENT-Montréal, deciduous species tended to occupy higher strata of the canopy than coniferous species (Urgoiti et al. 2023a). The species in these groups vary in both height at the crown top (due to variation in productivity) and at the crown base (due to variation in both productivity and self-pruning strategies). When pairing deciduous and evergreen species, deciduous species are likely to have benefited from competitive relaxation, and accordingly constructed much larger crowns (Fig. 4). These deciduous species likely have better light-use efficiency under high light, and their sparser crowns allow a relatively high fraction of light to filter down to less dominant evergreen species. Most of the evergreen species appear to have been able to continue growing under this partial shade, albeit not always as quickly, using much of the light that was not intercepted by the more dominant deciduous species. As a result, the influence of competitive imbalance may not have been particularly severe. Indeed, shade-tolerant species may even be facilitated by large neighbors that shield them from harsh microclimates (Kothari et al. 2021).

Many studies have found evidence for a trade-off between growth (or photosynthetic physiology) under high light and under low light (Pacala et al. 1994; Walters & Reich 2000; Baltzer & Thomas 2007a; Lusk & Jorgensen 2013; Falster et al. 2018). The success of mixtures after canopy closure may depend on an arrangement where the species that achieve early dominance are those that grow well under high light, and the species relegated to less dominant positions are those that can keep growing under relatively low light (Williams et al. 2021). This arrangement may be favored by the fact that rapid growth enables early dominance. However, if the dynamic were inverted—perhaps due to other factors such as dispersal timing—and more shade-tolerant species achieved early dominance, the more shade-intolerant species may not be able to persist beneath. Even without this sort of inversion, extreme competitive imbalance may weaken diversity-productivity relationships (Yachi & Loreau 2007), and there is some evidence that this may indeed occur in late succession (Yi et al. 2022).

Further investigation of tree diversity, crown architecture, and ecosystem function will likely benefit from the growing accessibility of technologies like above-canopy LiDAR and terrestrial laser scanning (TLS). For example, although we show that total crown volume increases with diversity in self-pruning strategies, we cannot state with certainty that total leaf area increases without having measured leaf area density. LiDAR and TLS offer the potential to map full leaf area profiles and segment individual trees (Lines et al. 2022; Owen & Lines 2024), which would enable studies of interactions between tree crowns in unprecedented detail.

In conclusion, we present the first major comparative study of self-pruning among trees in a light-limited forest. We find that species’ variation in self-pruning strategies can be explained by major axes of functional variation, including shade tolerance and the plant economic spectrum. These patterns are consistent with the idea that the light threshold of self-pruning (which we call *L_base_*) is determined by the amount of shading required for the branch to become carbon-depleted. Alongside the strong interspecific variation in *L_base_*, we also found distinctive patterns of plasticity: while acquisitive species self-pruned under deeper shade around acquisitive neighbors, conservative species self-pruned under deeper shade around conservative neighbors. Broader comparative studies could test the generality of these patterns and uncover further nuances in trees’ functional strategies. At the plot level, higher levels of diversity in *L_base_* correspond with greater basal area and canopy packing. Moreover, the relationship with canopy packing is strengthened by intraspecific adjustments in crown depth between monocultures and mixtures, which allows more acquisitive species to benefit from competitive release when growing with more conservative species. These findings reinforce the strong links between diversity in canopy architecture and productivity in young, light-limited forests. We propose that *L_base_* may serve as a simple crown-level functional trait to explain how the interactions between individual trees competing for light shape the structure and development of forest stands.

## Acknowledgements

UQAM and IDENT-Montréal are located on unceded Indigenous land. This land is recognized as the territory of the Kanien’kehà:ka Nation and has long served as a place of meeting and exchange for many other Indigenous nations. We owe our deep thanks to the many researchers and interns who have studied and maintained the IDENT-Montréal experiment during the first nine years of its existence. McGill University provided access to the land on which IDENT-Montréal sits. Peter Reich contributed to the initial design of the experiment. Daniel Lesieur and Mélanie Desrochers from the Centre d’étude de la forêt provided much valuable help with data management and GIS for positioning trees and plots. Eric Searle provided valuable advice on statistical analyses. Maria Faticov, Sarah Tardif, Charlotte Langlois, Davia Yahia, Elyssa Cameron, and other members of the Paquette Lab provided useful feedback on an earlier draft. AP received funding from the Fonds de recherche du Québec—Nature et technologies (FRQNT; #267091) and the Natural Sciences and Engineering Research Council of Canada (RGPIN-2018-05201). WSK received funding from the McIntire-Stennis Forest Research Program of the United States Department of Agriculture. CM received funding from a Canada Research Chair in the Resilience of Forests to Global Change.

## Data availability

All data collected as part of the self-pruning survey, as well as neighborhood- and plot-level variables needed to reproduce analyses, will be archived on a data repository in the very near future, or (at the latest) upon acceptance for publication. Original IDENT inventory data will be shared upon request, contingent on acceptance of IDENT’s terms and conditions. All original code used in data processing and analysis will be archived on Zenodo prior to publication.

## Author contributions

AP and CM designed the IDENT-Montréal experiment and funded this study. JU, AP, CM, and WSK conceived the project with substantial later contributions by SK. JU carried out field work and initial data curation with supervision from AP, CM, and WSK. SK performed statistical analyses and wrote the first draft of the manuscript with contributions from JU and AP; all authors contributed to revisions.

## Supporting Information

### Crown volume estimation and canopy packing

We estimated crown volumes based on crown depth (*D*), maximum crown radius (*R_max_*), and a shape parameter (*β*), borrowing a stylized representation of tree crowns from Purves et al. (2007) and Jucker et al. (2015). Crown depth can be defined as *D* = *T – B*, where *T* is the height of the crown top and *B* is the height of the crown base. In this notation, the radius *R* at any given height *h* between *B* and *T* can be described as:

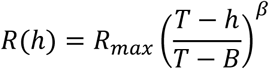

This function takes on the value of *R_max_* at *h = B* and 0 at *h = T*. The stylized crown shape results from its solid of revolution around the line *R* = 0. As derived by Jucker et al. (2015) using the shell method, the solid of revolution has the volume:

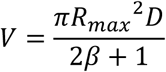

We used this formula to estimate the volume of each crown in the self-pruning survey.

The parameter *β* determines the curvature of the crown: for example, *β* = 0 results in a cylinder, *β* = 0.5 results in a paraboloid, and *β* = 1 results in a cone. Values of *β* were determined at the species-level based on the appendices to Purves et al. (2007). While it would have been preferable to have individual-level estimates of crown shape as in Williams et al. (2017), such measurements are extremely labor-intensive. As Jucker et al. (2015) show, the resulting estimates of crown volume are much less sensitive to *β* than to crown radius and crown depth. In this study, having the minimum species *β* value would only result in 13.9% more volume than the maximum value, holding crown depth and radius equal. In a broader sense, our reliance on Purves et al. (2007)’s tractable but rather constrained representation of crown geometry may somewhat bias our estimates of total canopy volume (Owen & Lines 2024). We acknowledge this limitation but suggest that there is no obvious reason to believe that such bias would influence comparisons among plots.

We considered three approaches to representing living crowns of a given species in a given plot. In the first approach, which we call the “actual crowns approach,” we used the living trees of that species in that plot measured in the self-pruning survey and estimated the volume of each one. In the second approach, called the “no-plasticity approach,” we aimed to assess the impact of self-pruning by evaluating whether we would observe more or less canopy packing in mixtures if trees did not adjust *D* between monocultures and mixtures. To this end, we sampled *R_max_* from living trees of the species in the plot and (for each mixture plot) sampled *D* from the monoculture of the same species in the same block. We performed 500 sampling iterations to simulate 500 crowns of each species in each plot.

Since there is a weakly positive empirical relationship between *R_max_* and *D*, the procedure of randomly assigning *R_max_* to *D* induces a small downward bias in estimates of crown volume. (To get an intuition for this idea, consider that the summed area of two rectangles each with side lengths *x* and *2x* is less than the summed areas of a square with side length *x* and a square with side length *2x*.) In our third approach, which we call the “independent draws with plasticity approach,” we sought to evaluate whether this bias could have altered our inferences of how self-pruning influences canopy packing. To this end, we also performed simulations where *D*, *R_max_*, and mortality rates were all drawn from the values in the same plot, but where *D* and *R_max_* for calculations of crown volume were independently sampled at random from trees in that plot in the self-pruning survey. As in the no-plasticity approach, we simulated 500 crowns of each species in each plot.

Comparing estimates of canopy packing between the actual crowns approach and the independent draws with plasticity approach confirmed the existence of a slight negative bias in approaches that involve drawing crown radius and depth independently. The magnitude of this bias was much smaller than the difference between the actual crowns approach and the independent draws without plasticity approach in mixture plots (Fig. S5), indicating that this difference was not an artifact of the bias. Nevertheless, to avoid overstating the difference between canopy packing with and without plasticity in *D*, we primarily compare the independent draws approaches with and without plasticity, since both are influenced by the same bias.

Regardless of the approach for estimating crown volumes of living trees in each species and plot, we aimed to scale these values up to total crown volume per unit area in each plot. Here, we only considered the inner 6 × 6 trees (9 m^2^), leaving out edges. Since our self-pruning survey only included living trees, we used the growth survey data to calculate the fraction of living trees of each species in each plot in 2018. We then estimated crown volume per unit area (*V*, in m^3^ m^-2^) as:

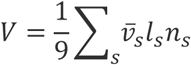

where ν̅_s_ is the mean crown volume of measured (living) trees of species *s* in the plot, *l_s_* is the fraction of trees of species *s* in the plot that were still alive, and *n_s_* is the number that were planted at the start of the experiment.

In statistical analyses of crown volume, we left out three plots where a tree species had at least one living individual in the plot, but the species was not surveyed in the self-pruning survey. In two cases, this was due to an error in following survey procedures, while in the remaining case the non-surveyed species had been planted by mistake.

### Net biodiversity effect on productivity and canopy packing

In many studies of the relationships between biodiversity and ecosystem functions, variation due to species composition makes it hard to isolate the influence of diversity on ecosystem function. As a result, researchers often calculate the net biodiversity effect (NBE), which quantifies the extent to which ecosystem function in mixtures deviates from expectations based on monocultures (Trenbath 1974).

In the main text, we assess the relationship between plot basal area (as a measure of productivity) and each of three variables quantifying functional diversity: the diversity of functional traits, the diversity of shade tolerance, and the diversity of *L_base_* (Fig. 5). Here, we report results from analyses that, rather than simply using basal area, use the NBE on basal area, quantifying how much productivity exceeds monoculture-based expectations.

For all mixtures of native species in all four blocks, we calculated the NBE on basal area as:

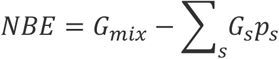

where *G_mix_* is the basal area of a given plot, *G_s_* is the basal area of a monoculture of species *s* in the same block, and *p_s_* is the proportion of species *s* planted in the mixture plot. Diversity in *L_base_* was better correlated with the NBE (*t*(98) = 5.04, *R^2^* = 0.206, *p* < 0.001; Fig. S4) than diversity in functional traits (*t*(98) = 3.61, *R^2^* = 0.117, *p* < 0.001) or diversity in shade tolerance (*t*(98) = 3.68, *R^2^* = 0.122, *p* < 0.001).

As with productivity, we found a positive relationship between total crown volume and diversity in functional traits or *L_base_*. However, for the same reason as in our analyses of productivity, we also calculated the NBE in crown volume in each mixture plot as:

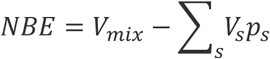

where *V_mix_* is the estimated total crown volume of the mixture plot, *V_s_* is the estimated total crown volume of a monoculture of species *s* in the same block, and *p_s_* is the proportion of species *s* planted in the mixture. Based on the independent draws with plasticity approach, the NBE on crown volume increased with both diversity of functional traits (*t*(43) = 4.84, *R^2^* = 0.353, *p* < 0.001; Fig. S5) and of *L_base_* (*t*(43) = 4.45, *R^2^* = 0.315, *p* < 0.001). On average, there was a positive NBE of canopy packing in both mixtures simulated with plasticity (one-sample *t*-test; average: 2.32 m, *t*(44) = 5.45, *p* < 0.001) and without, to a lesser degree (average: 1.45 m, *t*(44) = 4.96, *p* < 0.001).

**Fig. S1:**
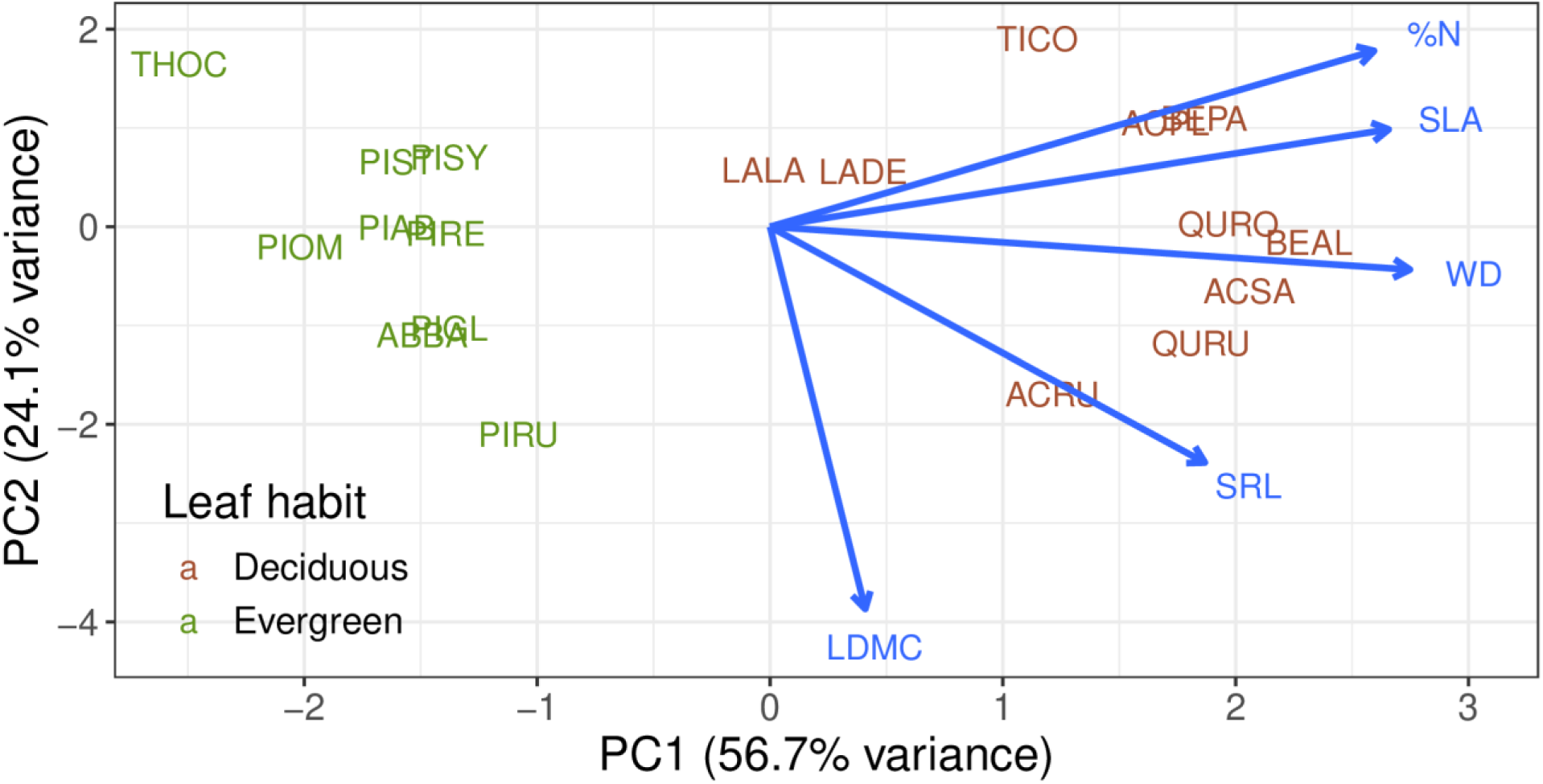
A biplot of species in trait space following principal components analysis on the trait correlation matrix. Arrows represent loadings. The PCA includes the twelve native species in the experiment (listed in Table 1) as well as seven exotic species that were planted in the experiment, but not considered as focal species here: *Acer platanoides* L. (ACPL), *Larix decidua* Mill. (LADE), *Picea abies* (L.) H. Karst. (PIAB), *Picea omorika* (Pančić) Purk. (PIOM), *Pinus sylvestris* L. (PISY), *Quercus robur* L. (QURO), and *Tilia cordata* Mill. (TICO).

**Fig. S2:**
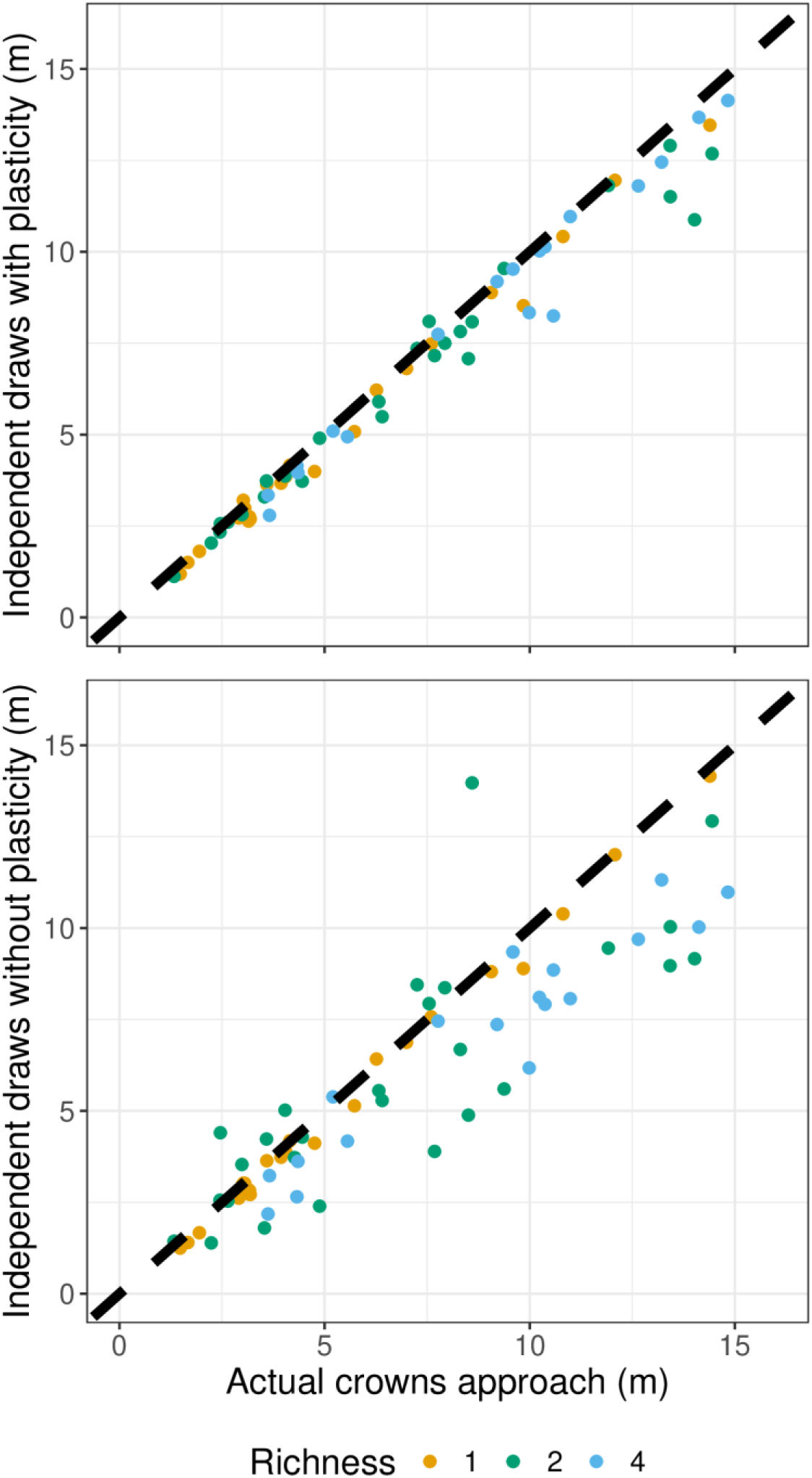
Estimates of crown volume per unit ground area, comparing the actual crowns approach with the two simulations: the independent draws without plasticity approach (top), and the independent draws with plasticity approach (bottom). The thick dashed line is the 1:1 line.

**Fig. S3:**
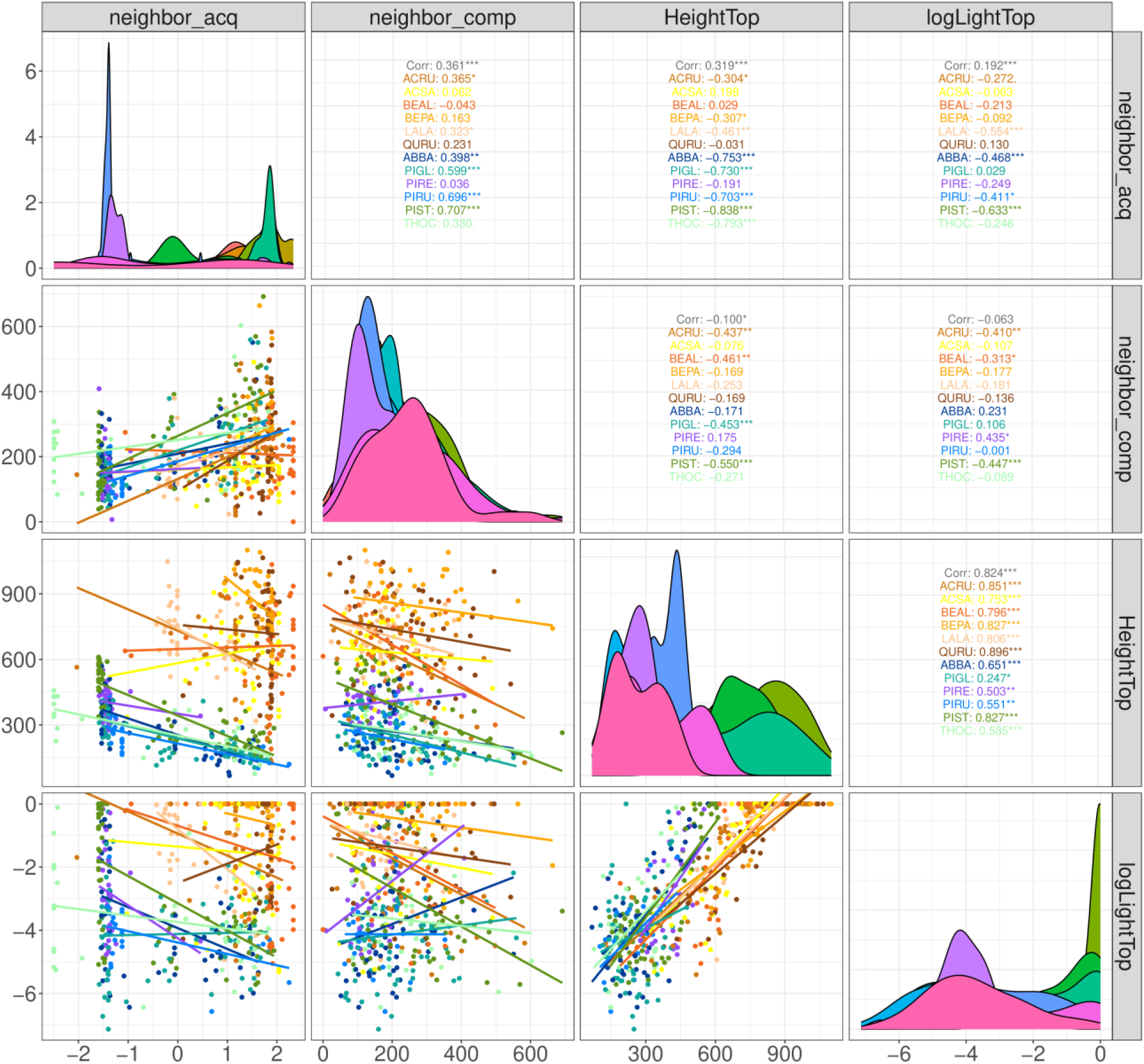
Pairwise correlations among neighborhood variables: height of focal individual, top light of focal individual, NCI, and neighborhood acquisitiveness.

**Fig. S4:**
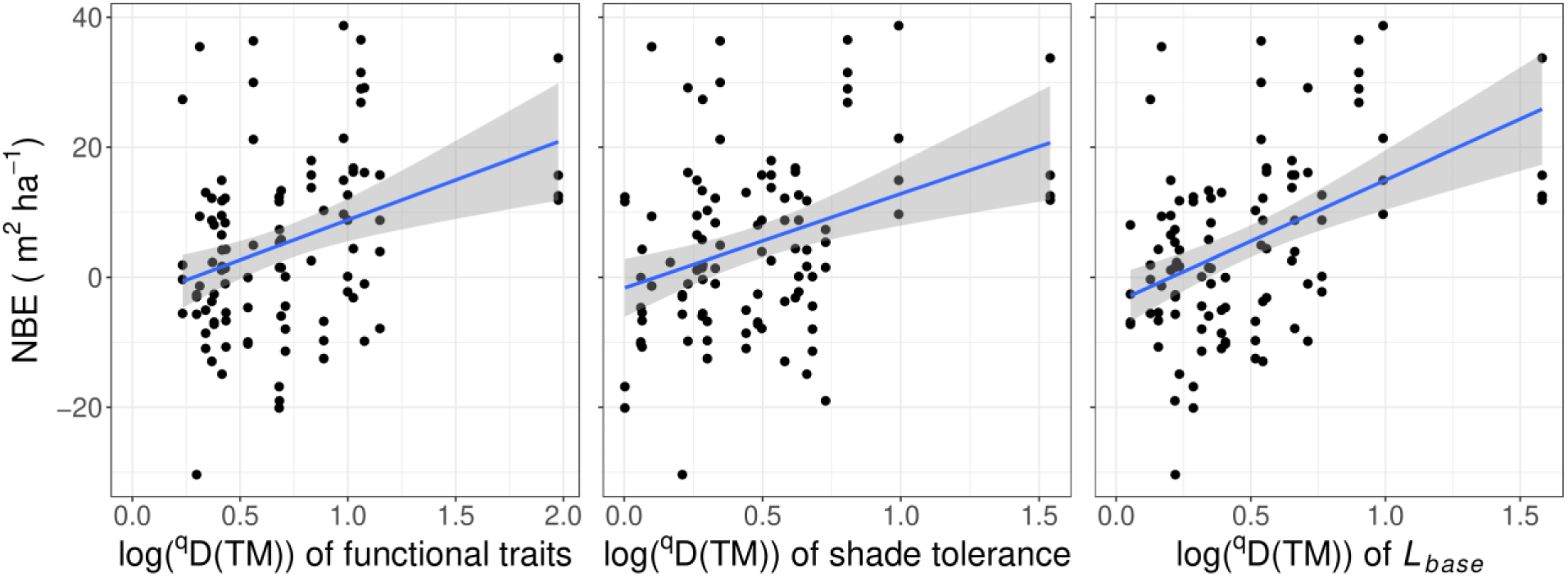
Correlations between the net biodiversity effect (NBE) on basal area and diversity of functional traits (left), shade tolerance (middle), and *L_base_* (right) among mixture plots. Blue lines are OLS regression lines, with 95% confidence intervals shaded in gray.

**Fig. S5:**
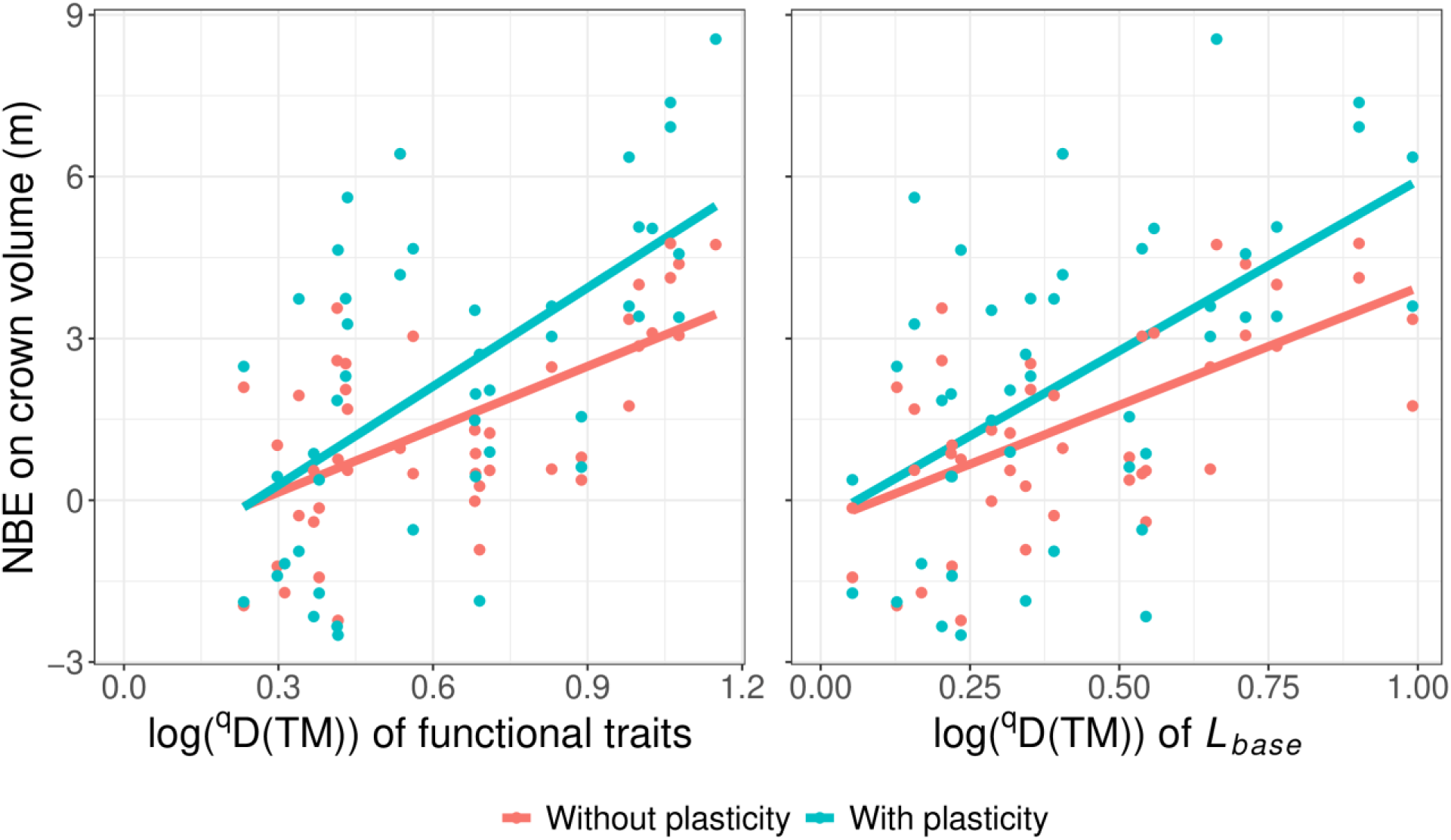
The relationship between the net biodiversity effect (NBE) on total crown volume per unit ground area (m^3^ m^-2^ or m) and diversity in functional traits (left) or *L_base_* (right) in two sets of simulations: one where plasticity in crown depth between monoculture and mixtures is incorporated, and one where it is not. Lines represent OLS regression fits.

**Fig. S6:**
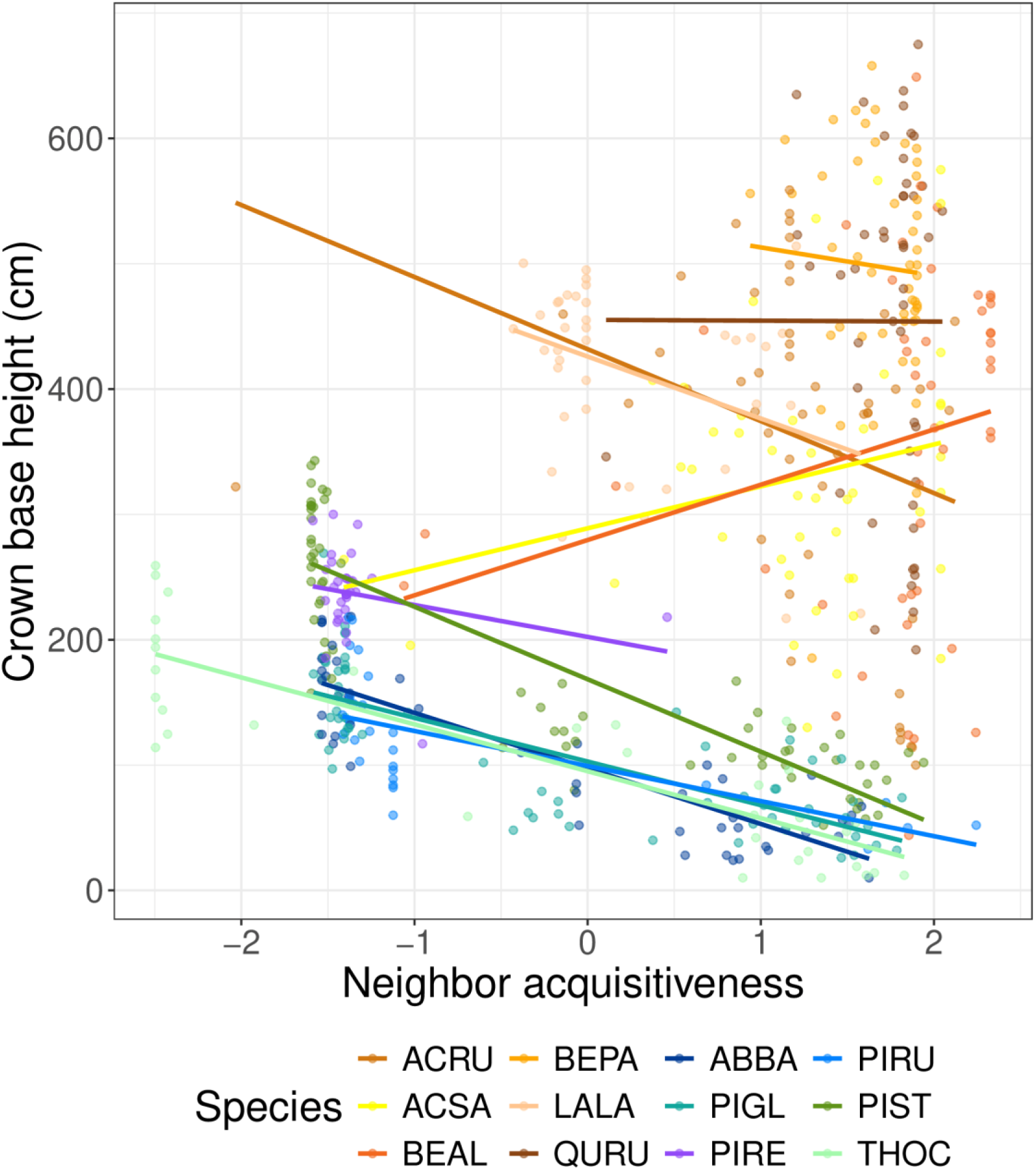
Relationships between neighbor acquisitiveness and crown base height across species. The best-fit lines are derived from species-specific OLS regressions.

